# Accurate human genome analysis with Element Avidity sequencing

**DOI:** 10.1101/2023.08.11.553043

**Authors:** Andrew Carroll, Alexey Kolesnikov, Daniel E. Cook, Lucas Brambrink, Kelly N. Wiseman, Sophie M. Billings, Semyon Kruglyak, Bryan R. Lajoie, June Zhao, Shawn E. Levy, Cory Y. McLean, Kishwar Shafin, Maria Nattestad, Pi-Chuan Chang

## Abstract

We investigate the new sequencing technology Avidity from Element Biosciences. We show that Element whole genome sequencing achieves higher mapping and variant calling accuracy compared to Illumina sequencing at the same coverage, with larger differences at lower coverages (20x-30x). We quantify base error rates of Element reads, finding lower error rates, especially in homopolymer and tandem repeat regions. We use Element’s ability to generate paired end sequencing with longer insert sizes than typical short–read sequencing. We show that longer insert sizes result in even higher accuracy, with long insert Element sequencing giving noticeably more accurate genome analyses at all coverages.

## Introduction

Sequencing the genomes and transcriptomes of organisms enables diagnosis of genetic diseases^1–3^, discovery of gene-trait associations^4^ for drug discovery^5^ and agriculture^6^, creation of reference genomes^7^, resources for genetic variant annotation^8^, and imputation methods^9^.

Initially, efforts to assess sequencing accuracy used indirect factors, such as the ratio of transition to transversion in variant calls or concordance with Mendelian inheritance^10^. The ability to assess accuracy was transformed by the Genome in a Bottle standards, a set of 7 human cell lines whose genomes were extensively characterized with multiple technologies, analysis methods, and manual curation^11–15^. This resource, combined with community competitions^16^ and comparisons^17^ expanded the ability to detect accuracy improvements beyond the accuracy of current individual methods, which allowed a burst of innovation in both sequencing^18^ and analysis^19–21^ methods to demonstrate the validity of their innovations.

A new sequencing method based on sequencing by avidity rather than sequencing by synthesis developed by Element Biosciences can generate short-read sequencing at high yield and efficient cost, along with per-base accuracies that reportedly exceed Illumina sequencing^22^. Because the reported metrics focus on the accuracy of individual reads this work assesses how the read level accuracies of Element correspond to accuracy of full genome sequencing including both mapping and variant calling of the Genome in a Bottle samples.

We observe that Element sequencing enables noticeably higher accuracy across a range of coverages from 20x-50x. The increase in accuracy was most notable at lower coverages (20x-30x). We identified genome contexts where Element had substantial improvements in accuracy, specifically tandem repeats and homopolymers, with a noticeable reduction in read soft-clipping due to loss of quality later in the read at these contexts.

One new property of Element’s AVITI platform is the ability to generate paired-end sequencing data with longer insert sizes (the distance between the paired reads) than is typical with Illumina preparations. By investigating Element sequencing runs performed with libraries that had longer insert size distribution (with a template length of >1000 base pairs as opposed to 350-500 base pairs), we identified a strong positive effect with longer insert sizes for Element sequencing. The long insert Element sequencing outperformed both Illumina and standard insert Element sequencing at each coverage threshold.

### Comparing variant calling accuracy

We compared genome analysis accuracy between Illumina and Element sequencing in typical use cases. Individual sequencing runs of high sequencing coverage of Illumina^23^ and Element were downsampled to equal number of starting reads for 20x, 30x, 40x, and 50x coverage. These reads were mapped with BWA MEM^24^ to the GRCh38 reference^25^. Sequencing runs from HG001, HG002, HG003, and HG005 were analyzed from both technologies. Variants were called with DeepVariant v1.5^26^, which has been jointly trained with both Illumina and Element data in the single release model. All comparisons between technologies use this same DeepVariant model. HG003 is withheld from training all DeepVariant models and that sample is used for whole genome holdout test datasets. Chromosome 20 is withheld from training in all samples and is used for comparison on other samples.

To assess accuracy, we used Hap.py^13^ to compare the resulting VCF against the v4.2.1 Genome in a Bottle truth set^15^ used in the PrecisionFDA v2 Truth Challenge^16^. Element sequencing had noticeably higher accuracy (both precision and recall) compared to Illumina at the 20x coverage point, but the difference narrowed at higher coverage **(Figure 1A, Supplementary Figure 1)**.

**Figure 1.**
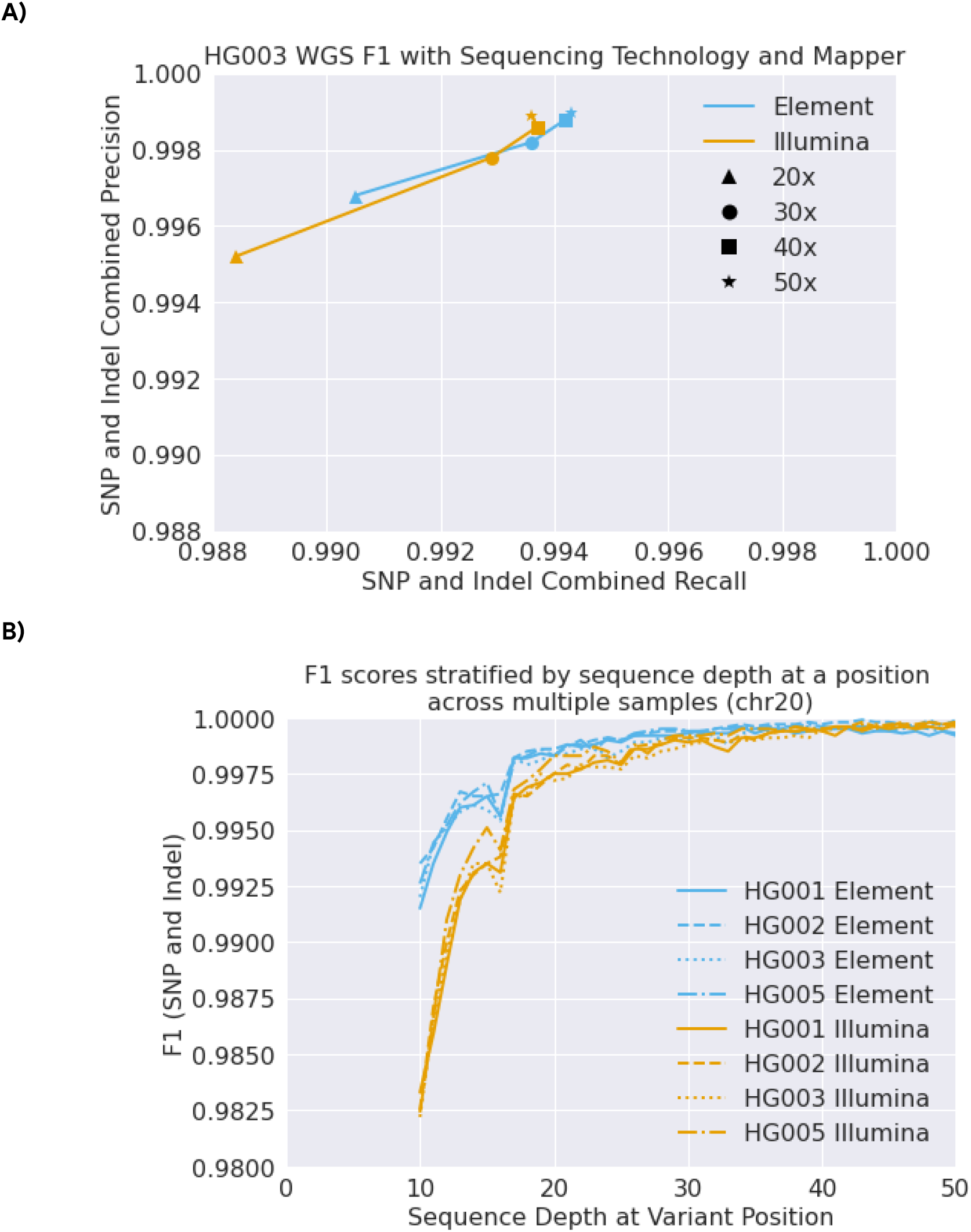
Variant calling accuracy for Element and Illumina sequencing. **(A)** Variant calling accuracy (precision and recall) with 20x-50x sequencing depth assessed by Genome in a Bottle. **(B)** Accuracy for Element and Illumina when matching coverage per position between the technologies.

A sample with 30x average sequencing coverage will have a distribution of coverages across the genome, for example some positions will be covered at 20x and others at 40x. To look more directly at the effect of coverage on accuracy for Illumina and Element, we downsampled at 1x intervals from 50x to 10x (a total of 40 variant calling runs per sample) across chromosome 20, which is always withheld from DeepVariant training in all samples. We collected the hap.py results for all variant call files, aggregated all calls, and stratified these calls by the sequencing depth at a given position. This allowed us to assess performance on coverage-matched positions across a large coverage range. This revealed larger differences in accuracy at lower coverages between Element and Illumina **(Figure 1B)**. Element had a clear advantage in the 30x-40x coverage range as well.

The first step of DeepVariant’s variant calling method uses a heuristic process, conceptually similar to Samtools^27^ bcftools or GATK^28^ which uses observed allele frequencies to propose positions as candidate variants. In the second stage, a convolutional neural network either rejects these candidates as false, or determines they are true and assigns their genotype. In order for a false candidate to be generated, at least two reads must support the candidate and at least 12% of total reads for SNPs or 6% for Indels. At the candidate variant level, we noticed more substantial differences between Element and Illumina runs. Element runs had fewer rejected candidates (**Figure 2A)**. The observation of lower false candidate generation was consistent with the reported higher overall accuracy of Element reads^22^. However, the magnitude of the difference seen here should require concentration of errors in certain contexts, as a 12% support rate is much higher than overall Illumina sequencing errors rates.

**Figure 2.**
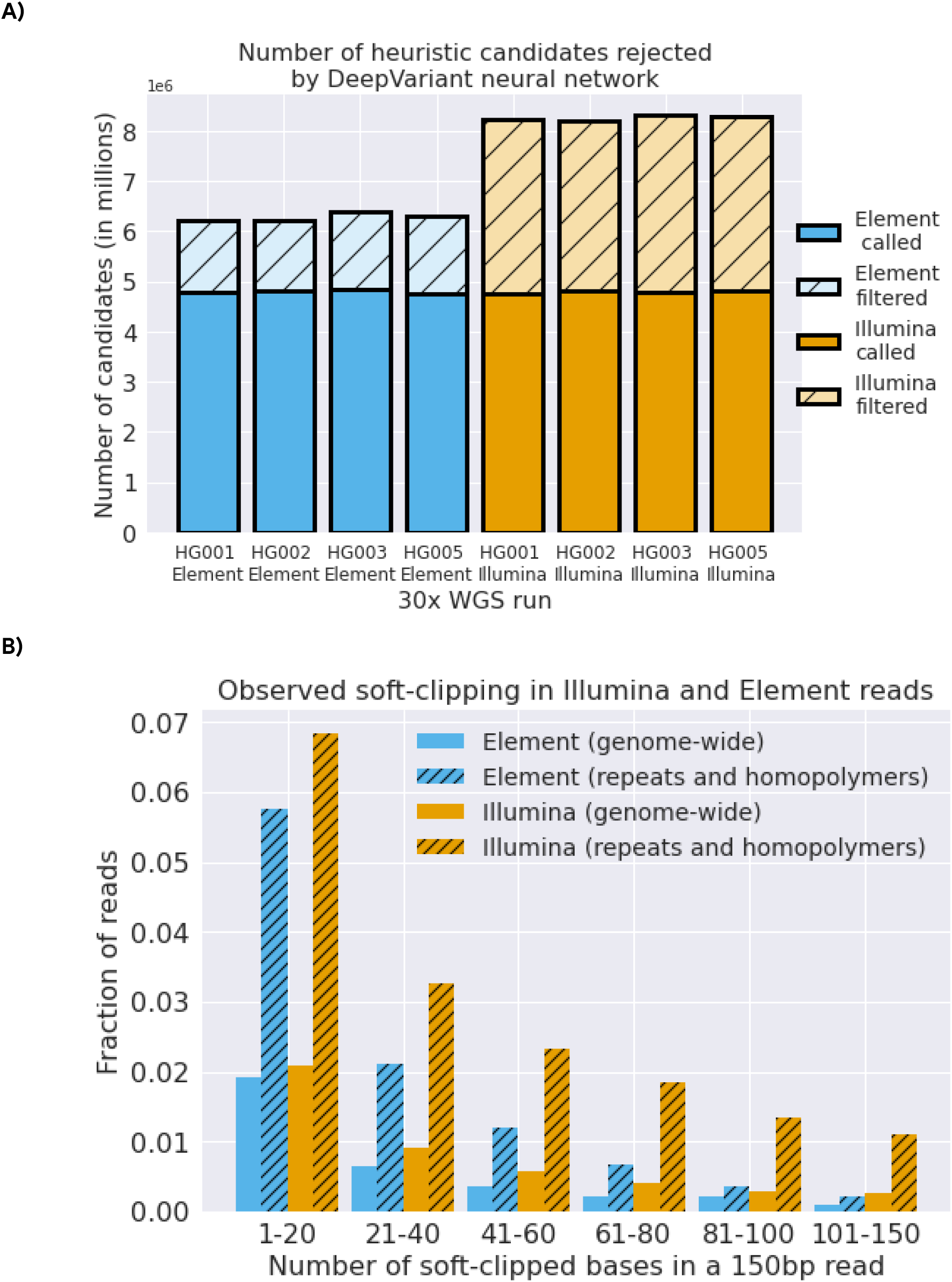
Heuristic approaches which depend on read error rates indicate higher accuracies in Element data. **(A)** Proportion of candidates proposed by heuristic logic which are rejected as false positives by DeepVariant’s neural net. **(B)** Proportion of soft-clipped bases in Element and Illumina reads across the genome and in repetitive regions.

The large differences in candidate generation rates between Illumina and Element seemed unlikely to be fully explained by random error rates. Instead, reads going out of sequencing phase in regions difficult to resolve by sequencing by synthesis could generate the error rates required to make candidates. In sequencing by synthesis, reads go out of phase when they hit certain contexts (e.g. homopolymers and tandem repeat runs) that break up the synchronous replication of the cluster, so individual molecules are replicating at different parts of their template^29^. This degrades the sequencing quality and produces errors.

The read mapping step can occasionally identify that a read has useless sequence after a certain point and soft-mask the read. We observed a higher proportion of soft-masked bases in Illumina, which was much more pronounced in repeat regions and homopolymers from the Genome in a Bottle stratifications^30^ (**Figure 2B)**. This is consistent with Element having a noticeable improvement in read phasing over difficult contexts.

### Investigating base-level concordance through T2T assemblies

Recently, a highly accurate. telomere-to-telomere (T2T) assembly of chromosome Y was completed for HG002^31^. This allows us to investigate the empirical accuracy of Element sequencing at the base-level within reads by mapping HG002 and HG003. HG003 is the father of HG002 and transmits the same Y-chromosome to the T2T assembly.

We used the Bam Error Stats Tool (BEST)^32^, which was developed to quantify errors in sequencing technology at the read level by comparing reads to a reliable assembly. Reads at MAPQ60 were used to greatly reduce mapping bias. Consistent with the observations from variant calling, we observe empirical concordance of HG002 and HG003 is higher with Element samples than with Illumina samples. Mismatch rates were 2.4 to 3.3 fold higher in Illumina reads compared to Element **(Figure 3A)**. We also compare the predicted base quality values and find their calibration consistent between Element samples and well-calibrated beyond predicted Q40 (99.99% accuracy). **(Figure 3B)**.

**Figure 3.**
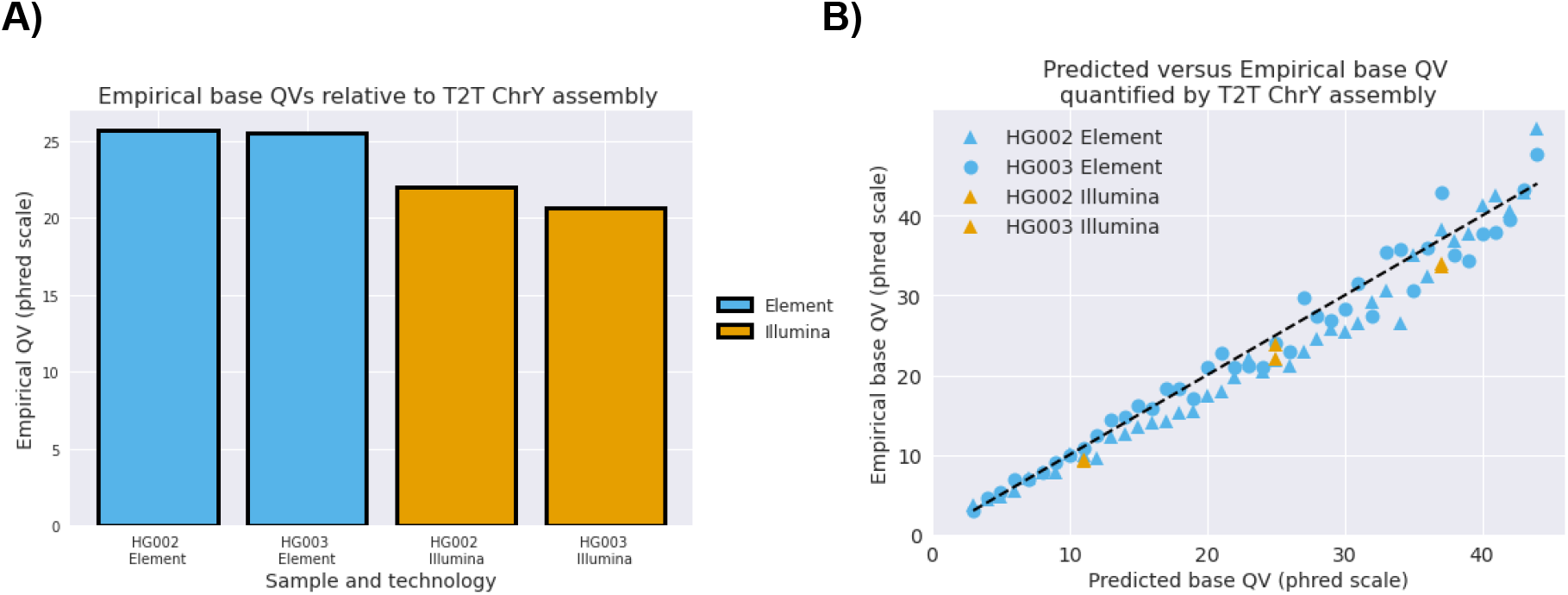
Read accuracy and calibration assessed by concordance with T2T assembly of chromosomeY. **(A)** Empirical QV for Element and Illumina sequencing runs for two samples (HG002 and HG003). Errors are averaged across all reads mapped to chrY with MAPQ60 in the T2T-XY 2.7 confident regions. **(B)** Calibration between sequencer-predicted base quality (x-axis) with empirical error rate (y-axis). The y=x line represents perfect calibration. The same MAPQ60 reads are used in both comparisons.

### Long insert sequencing improves genome analysis

Element has developed methods that allow libraries with insert sizes of >1000 base pairs as opposed to 350-500 base pairs to be sequenced efficiently. To test if longer inserts could improve Element sequencing accuracy, we received long insert sequencing runs with a median length of more than 1000bp (**Figure 4A)**. The same mapping and variant calling pipeline was used resulting in large improvements in recall **(Figure 4B, Supplementary Figure 2)**, suggesting that increasing insert length is a promising mechanism to increase variant calling comprehensiveness and accuracy in general.

**Figure 4.**
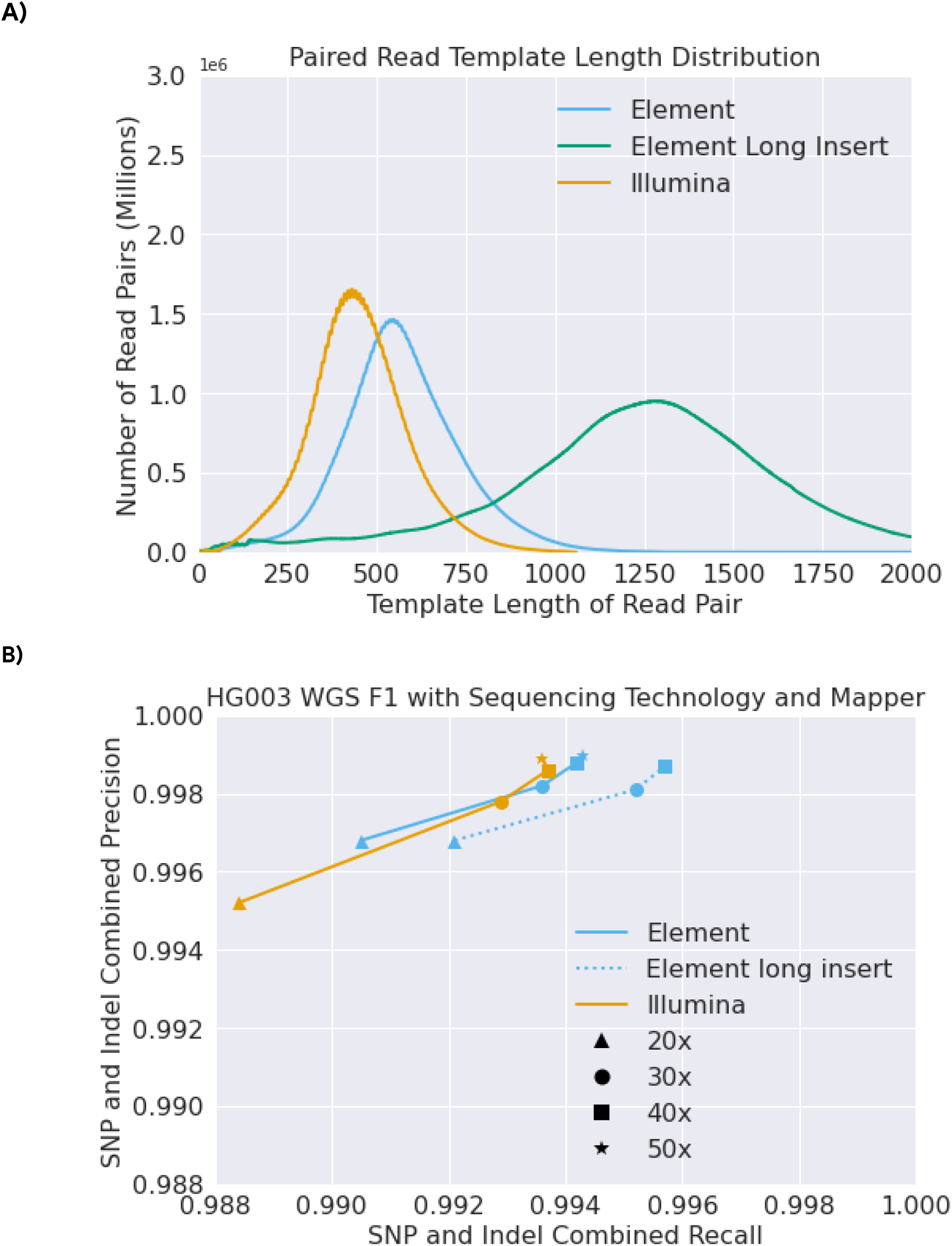
Longer sequencing inserts improve genome analysis accuracy for Element data. **(A)** distribution of template length for a long insert Element sequencing run versus standard insert Illumina and Element runs. **(B)** Variant calling accuracy (precision and recall) with 20x-50x sequencing depth assessed by Genome in a Bottle.

## Discussion

We have characterized the accuracy profile for analysis of human genomes with a new sequencing technology, Element AVITI that uses a sequencing by avidity approach rather than sequencing by synthesis. Element data achieves greater variant calling accuracy over a range of coverages, with especially improved accuracy in the 20x-30x coverage range. We identify certain sequence contexts in which Element outperforms Illumina reads, including in tandem repeats and homopolymers, as measured by soft-clipping rates. Finally, we show a positive effect on the accuracy of whole genome sequencing pipelines when using longer inserts for sequencing.

Although these investigations focus on whole genome analysis for germline variation at coverage ranges typically used for variant discovery, there are several other applications for which base-level accuracy is of greater importance. These applications include somatic sequencing for detection of subclonal acquired variants, deep sequencing of cancers, or analysis of cell-free DNA. For this application, only a few sequence reads may contain a variant at low allele fraction, and the ability to determine whether those bases reflect a real variant or an error depends highly on sequence quality. Similarly, low-pass sequencing of samples followed by imputation could benefit more, which has recently been investigated with Element sequencing^33^.

The high accuracy of Element in homopolymers and repeats could provide a unique ability to improve genome assemblies and reference resources by polishing remaining errors in these contexts which are difficult both for Illumina as well as long-read methods like Pacific Biosciences and Oxford Nanopore.

One caveat in analyses of accuracy is that the current (v4.2.1) Genome in a Bottle benchmarks do not cover the entirety of the Genome, due to the difficulty in mapping certain parts of it. Accuracy across the full genome, including these parts not covered by Genome in a Bottle is likely lower. The longer insert size Element runs might be able to better access parts of the genome which can’t be measured by these benchmarks, and could be another method to help expand the confident regions in future releases.

## Methods

### Data Availability

Illumina sequencing was taken from PCR-Free NovaSeq6000 data generated as described in Baid et al 2020^23^. Element sequencing data for whole genome comparison on HG003 was taken from the Cloudbreak release. Sequencing data from other samples was taken from earlier Element chemistries and made available by Element from: https://www.elementbiosciences.com/resources

FASTQ, BAM, VCFs, and analysis files are hosted publicly and available with no egress charge at: https://console.cloud.google.com/storage/browser/brain-genomics-public/research/element/

Accessible from GCP console at: gs://brain-genomics-public/research/element/

All contents of this folder are available via direct https links, an index of file urls can be downloaded at: https://storage.googleapis.com/brain-genomics-public/research/element/element_urls.txt

Within this folder there are five subfolders:

**candidates/** - VCF files for 30x sequencing of Illumina and Element multiple samples used to identify filtered candidates across samples (Figure 2A).

**chr20/** - Chromosome20 VCF files for 1x downsamples from 10x-50x of Illumina and Element samples. Used for Figure 1B.

**chry/** - ChromosomeY BAM, VCF, and Best analysis files used to assess read concordance with T2T assembly. Used for Figure 3A-B

**sequencing_files/** - Whole genome sequencing FASTQ files analyzed in this paper.

**wgs/** - FASTQ, BAM, VCF, and Hap.py files for HG003 Illumina and Element Cloudbreak, include standard (500bp) and long (1000bp) insert sizes. Used for figures 1A, 2B,4A-B.

### Protocol for long insert Element data

Covaris-sheared, PCR-free long insert libraries were prepared using the Kapa HyperPrep workflow. 1ug of HG002 and HG003 gDNA were mechanically sheared using the following Covaris program:

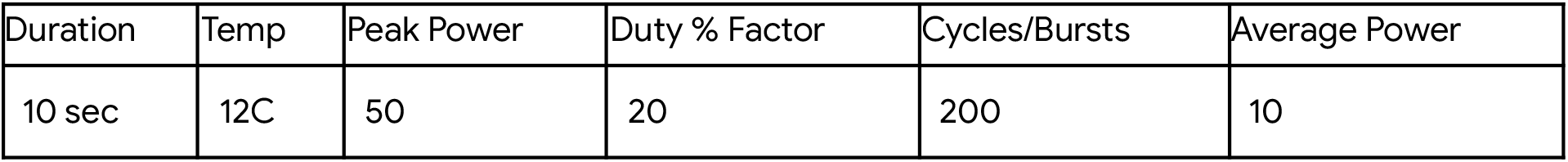

A narrow double-sided SPRI selection ratio of 0.3X/0.42X was used to select the long fragments. The Adept Rapid protocol was used for circularization. The libraries were sequenced on the Element AVITI system, 2x150 paired end reads with indexing, using a custom recipe for long inserts. The primary changes to the recipe involved increasing the amplification time to account for the increased insert length.

### Reference Genome used

GRCh38 with masking of certain false segmental duplications as recommended by Genome in a Bottle^34^ (GRCh38_masked_v2_decoy_excludes_GPRIN2_DUSP22_FANCD2.fasta.gz) was used for all germline variant calling pipelines. For BEST analysis ChrY of T2T-CHM13v2.0^31^ was used.

### Read Mapping

Mapping was performed with BWA v0.7.17 (r1188)^24^. Duplicate marking was performed with GATK v4.1.2^28^

~~~
BWA bwa mem -t ${THREADS} “${REF}” “${READS}” “${MATES}” -R
“@RG\\tID:${RGID}\\tPL:ILLUMINA\\tPU:NONE\\tLB:${RGID}\\tSM:${SAMPLE}
“ | samtools sort -O BAM -o ${BWA_OUTPUT}
MarkDuplicates java -jar gatk-package-4.1.2.0-local.jar
MarkDuplicates -I ${BWA_OUTPUT} -O ${OUTPUT} –M
${OUTPUT%.bam}.metrics
~~~

### Variant Calling

Variant calling was performed with DeepVariant v1.5 ^26^ using the WGS model.

~~~
sudo docker run \
 -v “${INPUT_DIR}”:”/input” \
 -v “${OUTPUT_DIR}:/output” \
 google/deepvariant:1.5.0 \
 /opt/deepvariant/bin/run_deepvariant \
 --model_type=WGS \
 --ref=/input/”${REF}” \
 --reads=/input/”${READS}” \
 --output_vcf=/output/”${READS}” \
 --num_shards=“$(nproc)” \
 --make_examples_extra_args=“downsample_fraction=${DFRAC}”
~~~

### Read and base level assessment

Assessment of base and read level accuracy was performed with BAM Error Stats Tool (BEST)^32^

./best ${BAM} GRCh38_masked_v2_decoy_excludes_GPRIN2_DUSP22_FANCD2.fasta ${PREFIX}

Assessment used only MAPQ 60 reads in the T2T-XY v2.7 confident regions. The confident BED file used is at:

https://storage.googleapis.com/brain-genomics-public/research/element/chry/T2T_chrY_confident.bed

### Variant accuracy evaluation

Accuracy evaluation was performed with hap.py^13^ using the v4.2.1^13,15^ truth sets from Genome in a Bottle.

~~~
sudo docker run -i \
 -v “${INPUT_DIR}”:”/input” \
 -v “${OUTPUT_DIR}”:”/output” \
 pkrusche/hap.py /opt/hap.py/bin/hap.py \
 /input/”${TRUTH_VCF}” \
 /output/${VCF} \
 -f /input/”${TRUTH_BED}” \
 -r /input/”${REF}” \
 -o /output/happy.output${PREFIX} \
 --engine=vcfeval
~~~

### Chromosome20 downsampling

To assess accuracy matching coverage at variant position for Figure 1B, downsamples at 1x intervals were conducted from 50x to 10x, and hap.py used to annotate variant calls as true positives, false positives, and false negatives. All calls for a given sample were aggregated over the files, and the sequence depth for each variant position was used to calculate total precision and recall.

### Read count and downsampling fraction for sequence data

The sequencing data used for this paper is available as FASTQ files at: https://console.cloud.google.com/storage/browser/brain-genomics-public/research/element/sequencing_files/

The number of reads for each sample (including both paired files) and the downsample fraction required to reach 30x coverage are:

| Sample | Number of Reads<br>(150bp) | Downsample fraction for<br>30x |
| --- | --- | --- |
| HG001 Novaseq | 1172815198 | 0.5115895505 |
| HG002 Novaseq | 1260451774 | 0.4760197989 |
| HG003 Novaseq | 1201016046 | 0.4995770057 |
| HG005 Novaseq | 1065206804 | 0.5632709045 |
| HG001 Element | 1162209576 | 0.5162580075 |
| HG002 Element | 2045150782 | 0.2933769017 |
| HG003 Element<br>(500bp insert) | 1554087230 | 0.7612767255 |
| HG003 Element<br>(1000bp insert) | 788149670 | 0.3860787145 |
| HG005 Element | 932153224 | 0.6436710023 |

## Competing interests

AC, AK, DEC, LB, CYM, KS, MN, and PC-C are employees of Google LLC and own Alphabet stock as part of the standard compensation package. KNW, SMB, SK, BRJ, JZ, and SEL are employees of Element Biosciences and hold stock options in the company.

## Supplementary Figures

**Supplementary Figure 1.**
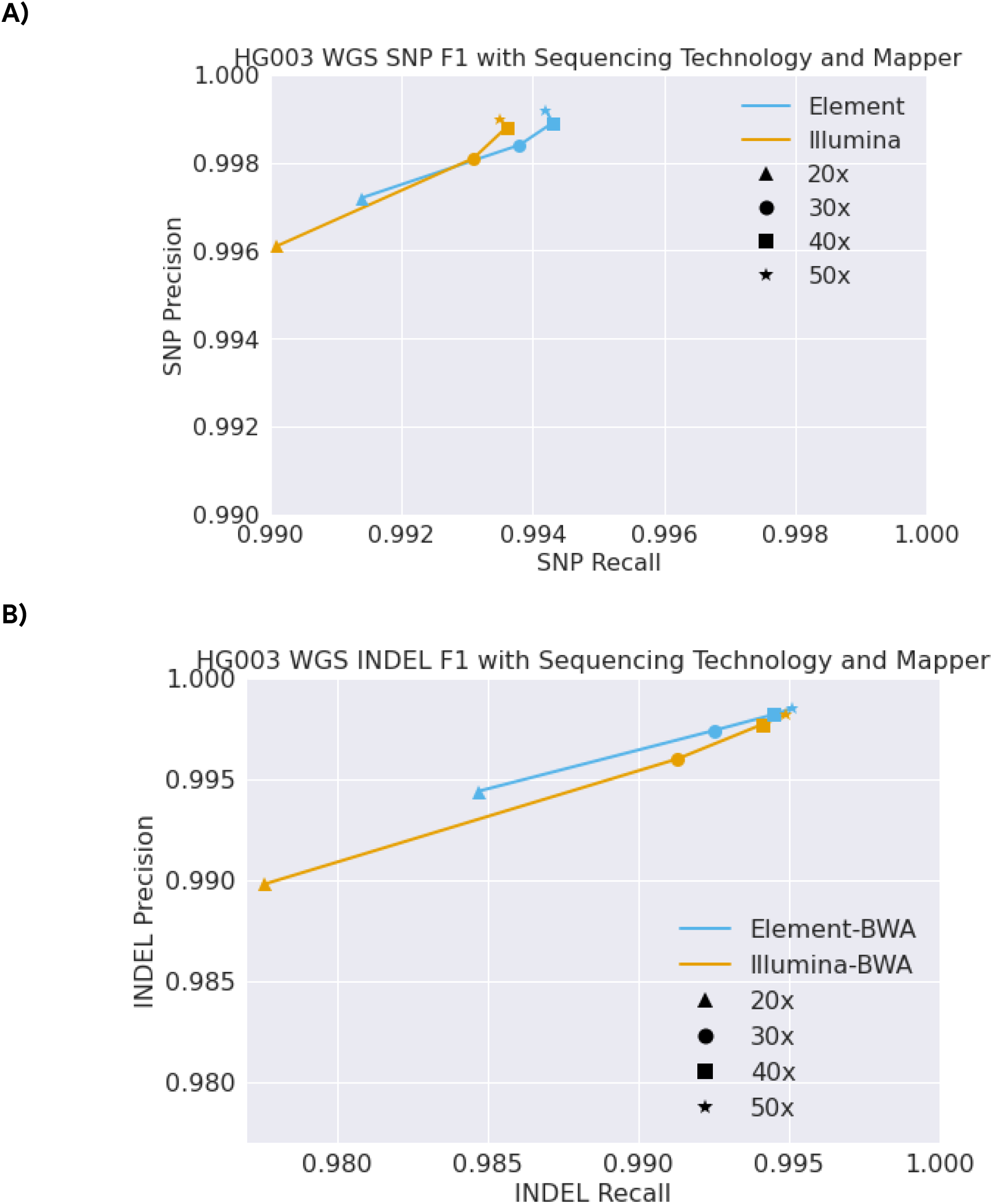
Variant calling accuracy for Element and Illumina sequencing separating SNP and Indels. **(A)** SNP Variant calling accuracy (precision and recall) with 20x-50x sequencing depth assessed by Genome in a Bottle. **(B)** Indel Variant calling accuracy (precision and recall) with 20x-50x sequencing depth assessed by Genome in a Bottle.

**Supplementary Figure 2.**
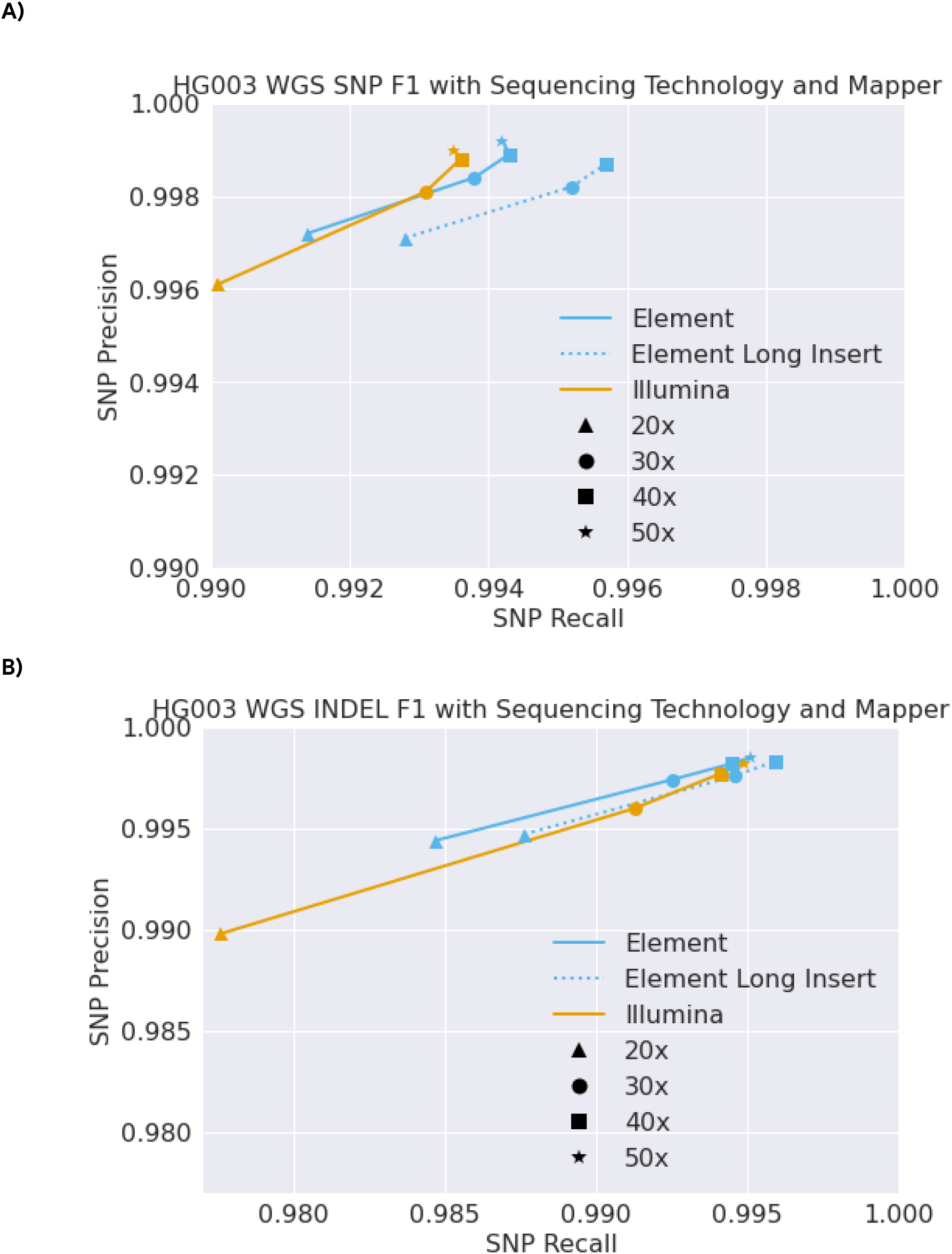
Longer sequencing inserts improve genome analysis accuracy for Element data separating SNPs and Indels. **(A)** SNP Variant calling accuracy (precision and recall) with 20x-50x sequencing depth assessed by Genome in a Bottle. **(B)** Indel Variant calling accuracy (precision and recall) with 20x-50x sequencing depth assessed by Genome in a Bottle.

## Notes

https://console.cloud.google.com/storage/browser/brain-genomics-public/research/element/

https://storage.googleapis.com/brain-genomics-public/research/element/element_urls.txt

## References

1. Gorzynski, J. E. et al.. Ultrarapid Nanopore Genome Sequencing in a Critical Care Setting. N. Engl. J. Med. 386, 700–702 (2022).

2. Saunders, C. J. et al.. Rapid whole-genome sequencing for genetic disease diagnosis in neonatal intensive care units. Sci. Transl. Med. 4, 154ra135 (2012).

3. AlDubayan, S. H. et al.. Detection of Pathogenic Variants With Germline Genetic Testing Using Deep Learning vs Standard Methods in Patients With Prostate Cancer and Melanoma. JAMA 324, 1957–1969 (2020).

4. Manolio, T. A. Genomewide association studies and assessment of the risk of disease. N. Engl. J. Med. 363, 166–176 (2010).

5. Peloso, G. M. et al.. Rare Protein-Truncating Variants in APOB, Lower Low-Density Lipoprotein Cholesterol, and Protection Against Coronary Heart Disease. Circ Genom Precis Med 12, e002376 (2019).

6. Snelling, W. M. et al.. Assessment of Imputation from Low-Pass Sequencing to Predict Merit of Beef Steers. Genes 11, p(2020).

7. Liao, W.-W. et al. A draft human pangenome reference. Nature 617, 312–324 (2023).

8. Karczewski, K. J. et al. The mutational constraint spectrum quantified from variation in 141,456 humans. Nature 581, 434–443 (2020).

9. Li, Y., Willer, C., Sanna, S. & Abecasis, G. Genotype imputation. Annu. Rev. Genomics Hum. Genet. 10, 387–406 (2009).

10. Eberle, M. A. et al. A reference data set of 5.4 million phased human variants validated by genetic inheritance from sequencing a three-generation 17-member pedigree. Genome Res. 27, 157–164 (2017).

11. Zook, J. M. et al. Integrating human sequence data sets provides a resource of benchmark SNP and indel genotype calls. Nat. Biotechnol. 32, 246–251 (2014).

12. Zook, J. M. et al. Extensive sequencing of seven human genomes to characterize benchmark reference materials. Scientific data vol. 3 160025 (2016).

13. Krusche, P. et al. Best practices for benchmarking germline small-variant calls in human genomes. Nat. Biotechnol. 37, 555–560 (2019).

14. Zook, J. M. et al. An open resource for accurately benchmarking small variant and reference calls. Nat. Biotechnol. 37, 561–566 (2019).

15. Wagner, J. et al. Benchmarking challenging small variants with linked and long reads. Cell Genom 2, (2022).

16. Olson, N. D. et al. PrecisionFDA Truth Challenge V2: Calling variants from short and long reads in difficult-to-map regions. Cell Genom 2, p(2022).

17. Foox, J. et al. Performance assessment of DNA sequencing platforms in the ABRF Next-Generation Sequencing Study. Nat. Biotechnol. 39, 1129–1140 (2021).

18. Wenger, A. M. et al. Accurate circular consensus long-read sequencing improves variant detection and assembly of a human genome. Nat. Biotechnol. 37, 1155–1162 (2019).

19. Sirén, J. et al. Pangenomics enables genotyping of known structural variants in 5202 diverse genomes. Science 374, abg8871 (2021).

20. Tetikol, H. S. et al. Pan-African genome demonstrates how population-specific genome graphs improve high-throughput sequencing data analysis. Nat. Commun. 13, 4384 (2022).

21. Scheffler, K. et al. Somatic small-variant calling methods in Illumina DRAGENTMSecondary Analysis. bioRxiv 2023–2003 (2023).

22. Arslan, S. et al. Sequencing by avidity enables high accuracy with low reagent consumption. Nat. Biotechnol. (2023) doi:10.1038/s41587-023-01750-7.

23. Baid, G. et al. An Extensive Sequence Dataset of Gold-Standard Samples for Benchmarking and Development. Cold Spring Harbor Laboratory 2020.12.11.422022 (2020) doi:10.1101/2020.12.11.422022.

24. Li, H. Aligning sequence reads, clone sequences and assembly contigs with BWA-MEM. arXiv [q-bio.GN] (2013).

25. Schneider, V. A. et al. Evaluation of GRCh38 and de novo haploid genome assemblies demonstrates the enduring quality of the reference assembly. Genome Res. 27, 849–864 (2017).

26. Poplin, R. et al. A universal SNP and small-indel variant caller using deep neural networks. Nat. Biotechnol. 36, 983–987 (2018).

27. Li, H. et al. The Sequence Alignment/Map format and SAMtools. Bioinformatics 25, 2078–2079 (2009).

28. DePristo, M. A. et al. A framework for variation discovery and genotyping using next-generation DNA sequencing data. Nat. Genet. 43, 491–498 (2011).

29. Pfeiffer, F. et al. Systematic evaluation of error rates and causes in short samples in next-generation sequencing. Sci. Rep. 8, 10950 (2018).

30. Dwarshuis, N. J. et al. StratoMod: Predicting sequencing and variant calling errors with interpretable machine learning. bioRxiv 2023–2001 (2023).

31. Rhie, A. et al. The complete sequence of a human Y chromosome. bioRxiv 2022–2012 (2022).

32. Liu, D. et al. Best: A Tool for Characterizing Sequencing Errors. bioRxiv 2022–2012 (2022).

33. Li, J. H. et al. Low-pass sequencing plus imputation using avidity sequencing displays comparable imputation accuracy to sequencing by synthesis while reducing duplicates. bioRxiv 2022–2012 (2022).

34. Behera, S. et al. FixItFelix: improving genomic analysis by fixing reference errors. Genome Biol. 24, 31 (2023).

